# Exploring the effects of Golgi Reassembly and Stacking Proteins in lipid membranes

**DOI:** 10.64898/2026.02.11.705364

**Authors:** Emanuel Kava, Leonel Malacrida, Marcela Díaz, Rosangela Itri, Antonio J. Costa-Filho

**Author notes:** To whom correspondence should be addressed: Antonio J. Costa-Filho, DF/FFCLRP/USP, Av. Bandeirantes 3900, Ribeirão Preto, SP, Brazil, 14040-901.

## Abstract

Golgi Reassembly and Stacking Proteins (GRASP) have been associated with Golgi-ribbon structure and unconventional vesicular protein secretion. In performing these functions, GRASPs consistently interact with membranes. The presence of lipid modifications, such as myristoylation, is a crucial consideration for obtaining detailed information about the interactions between GRASPs and membranes. Nonetheless, it has been overlooked in the literature so far. Here, we describe a reconstitution protocol for myristoylated human GRASP65 and GRASP55 in lipid model membranes, enabling investigation of their interactions using techniques ranging from structural characterization to spectroscopy and microscopy. Our results showed that myristoyl-anchored GRASPs can influence membrane dynamics, suggesting a possible role for their disordered SPR domain in this interaction.

## 1. Introduction

In the so-called conventional secretion pathway, the endoplasmic reticulum (ER) initiates protein synthesis and folding and begins glycosylation of a portion of the proteome^1^. Proteins synthesized in the ER are transported through coat protein complex II (COPII) vesicles to the Golgi, where post-translational modifications occur, such as phosphorylation and glycosylation^2^. The Golgi apparatus is essential for processing secretory and membrane proteins. It consists of laterally linked flattened cisternae, each characterized by distinct microenvironments across Golgi regions, where dynamic sorting of signaling proteins occurs. It is a highly polarized organelle with different enzymatic activities in its *cis*, medial, and *trans* regions^3^.

In mammalian cells, the maintenance of the Golgi structure was initially believed to involve two Golgi Reassembly and Stacking Proteins (GRASPs): GRASP65^4^ (gorasp 2) and GRASP55^5^ (gorasp 1), located in the *cis* and *trans*/medial Golgi, respectively. In 1997, Barr et al. reported the existence of an N-ethylmaleimide (NEM) sensitive protein with an apparent mass of 65 kDa (named GRASP65) and related to the Golgi stacking and reassembly upon mitosis. Additionally, GRASP65 is myristoylated at glycine 2 of its N-terminus and forms a complex with GM130, a golgin family protein implicated in vesicle docking^6^. Two years later, Shorter et al. identified a homologue of GRASP65 by observing a new 55 kDa protein recognized by one of the antibodies used against a conserved peptide of GRASP65 in different organisms. This representative, designated GRASP55, is also a NEM-sensitive component of the Golgi stacking machinery and interacts weakly with the C-terminus of GM130. Golgin45, the golgin partner of GRASP55, was reported by Short et al. as an essential complex for normal Golgi structure^7^.

Structurally, GRASPs are composed of two distinctly functional domains: the highly conserved N-terminal GRASP domain, which is composed of two PDZ subdomains in tandem, and the C-terminal serine and proline-rich (SPR) domain, which is intrinsically disordered and remarkably divergent even between GRASPs of the same organisms^8^. In humans, GRASPs are myristoylated at their N-termini, with this lipidation serving as an anchoring mechanism to the Golgi membranes, together with Golgin interactions^9^. Single and double knockouts of the human GRASP genes yielded controversial conclusions, casting doubt on the actual roles of these proteins in Golgi stacking^10–12^. Golgi organization remained unchanged in cells depleted of the gorasp1 and gorasp2 genes, provided that Golgin45 and GM130 were overexpressed^13^. Recently, two independent reports showed mammalian GRASPs were not essential for cell viability, and the absence of individual GRASPs did not significantly affect Golgi cisternal stacking^14,15^. According to these studies, GRASPs are dispensable for Golgi cisternal stacking but crucial for lateral connections that link stacks to form the Golgi ribbon^16^.

Recently, GRASPs were shown to have an essential role in the unconventional protein secretion (UPS) under cell stress^17–23^ with representatives of the GRASP family participating in the transport of leaderless cytosolic^20^ and membrane proteins^24,25^. In the case of human GRASPs, GRASP55 appears to be more involved in UPS, whereas GRASP65 has few reports of an active role in protein secretion ^26^. GRASP55 relocalization to the ER during stress^27^, and its co-localization with autophagosomes and late endosomes/lysosomes upon glucose deprivation^28^ have been reported. To be located in such regions, GRASP55 must leave the Golgi, where it is anchored via myristoylation and binding to Golgin45^29^.

Despite their frequent presence near or at membrane environments, the specific physicochemical determinants of GRASP interactions with membranes remain poorly understood. In this report, we aimed to use a previously established myristoylation protocol for the GRASP domain^30,31^ to modify full-length human GRASPs, thereby allowing us to investigate these determinants, in particular, the role of the SPR domain in the anchoring mechanism. To do so, we expressed and purified the myristoylated versions of the full-length and GRASP domains of GRASP65 and GRASP55 and reconstituted those lipidated proteins into lipid membrane vesicles. The protein-lipid interaction was then analyzed through spectroscopic methods, such as electron spin resonance (ESR), fluorescence, and microscopy methods, such as hyperspectral imaging (HSI).

## 2. Materials and Methods

### 2.1. Protein expression and myristoylation protocol

Human GRASPs and their GRASP domains sequences were inserted into a pET-28a(+) expression vector using XhoI and NcoI restriction enzymes, while the encoding sequence of N-myristoyltransferase 1 (NMT1) from *Candida albicans* was subcloned into a pET-22b expression vector using XhoI and NdeI enzymes. The two plasmids (GRASP– and NMT-containing) were transformed by electroporation into BL21 Star (DE3) competent cells and grown overnight in 20 mL of LB medium containing 1% agar, with kanamycin (50 μg/mL) and ampicillin (100 μg/mL). Single colonies were inoculated in 5 mL of liquid LB medium O.N. at 37 °C and 180 rpm of agitation, then transferred into a 2 L Erlenmeyer flask containing 1 L of the same medium, in the presence of antibiotics, 37°C and 180 rpm until reaching the optical density at 600 nm (OD600) between 0.4-0.6. Cells were induced using 500 μM of IPTG (Isopropyl β-D-1-thiogalactopyranoside), with the addition of 100 nM of myristic acid dissolved in pure ethanol, for 18 h at 18°C and 180 rpm.

### 2.2. Protein purification and fluorescent labelling

The purification of GRASP65, DGRASP65, GRASP55, DGRASP55 and myr-DGRASP55 was performed as previously described^30,32–34^. The following protocol was used to purify myr-GRASP65, myr-DGRASP65, and myr-GRASP55. Cells were harvested by centrifugation at 8,000×g for 5 minutes, then resuspended in 20 mL of buffer (HEPES 25 mM, NaCl 300 mM, β-DDM 0.03% w/v, pH 7.4) per liter of cell culture. The cells were lysed by sonication in a Branson 450 Digital Sonifier® (Branson Ultrasonics Corporation, Danbury, USA) with 8 min intervals of 15 s between 5 s ultrasound pulses. The lysate was clarified by centrifugation at 12,000×g for 20 min, and the supernatant was incubated with 4 mL of Ni^2+^ –NTA resin for 15 min, followed by washing with purification buffer (HEPES 25 mM, NaCl 150 mM, β-DDM 0.03% w/v, pH 8.0) containing 20 mM. The purified protein was eluted with 6 mL of the purification buffer containing 300 mM imidazole. The solution eluted from the nickel column was then concentrated in an Amicon Ultra-15 Centrifugal Filter (NMWL 10 kDa, Merck Millipore, Burlington, USA) up to a volume of 0.5 mL, and then injected into a size exclusion chromatography (SEC) column Superdex200 10/300 coupled to an *Äkta Purifier 10* FPLC system (GE Healthcare Life Sciences, Chicago, USA) using the purification buffer.

### 2.3. Circular Dichroism Spectroscopy

CD spectra were recorded using a Jasco J-810 spectropolarimeter (JASCO Corporation, Tokyo, Japan) equipped with a Peltier temperature controller. A quartz cell with a 1 mm optical path length was used as a sample holder. The scanning speed was 100 nm/min, the spectral bandwidth was 1.5 nm, the response time was 0.5 s, and the final spectrum was averaged over 9 accumulations. The protein concentration was 2 µM in a sodium phosphate buffer solution (20 mM, pH 7.4).

### 2.4. Protein reconstitution into lipid vesicles

The lipid 1,2-dipalmitoyl-sn-glycero-3-phosphocholine (DPPC) was solubilized in chloroform to a concentration of 35 mM. The appropriate volume of the lipid solution was then added to glass tubes and slowly dried under nitrogen. An additional drying step was performed in vacuum using a SpeedVac^TM^ concentrator (Thermo Electron Corporation, Milford, USA) for 2 hours to remove the residual chloroform. The lipid films were hydrated with protein solutions in a detergent-containing buffer, then incubated for 2 hours in an ultrasonic bath to solubilize any residual lipid film. The proteoliposomes mixed with detergent were then incubated with BioBeads at 4°C for 16 h in slow agitation. The liposome-reconstituted proteins were carefully separated from the adsorbent beads with a micropipette, followed by centrifugation at 18,000 × g for 40 min at 4°C to separate the lipid fraction containing the anchored protein. The sedimented proteoliposome was then resuspended in a detergent-free buffer (HEPES, 25 mM; NaCl, 150 mM; pH 8.0) for all subsequent experiments. The control lipid vesicles were prepared using the same procedure without protein.

### 2.5. Differential Scanning Calorimetry (DSC)

The effects of GRASPs on the phase transitions of DPPC vesicles were determined using the VP-DSC microcalorimeter from MicroCal (Microcal, Northampton, MA, USA). All samples were prepared using the reconstitution protocol. A heating rate of 1 °C/min over a temperature range of 10-60 °C was chosen. Before each scan, the samples were equilibrated at 10°C for 15 minutes. Manipulation and analysis of the thermograms, including subtraction of the buffer’s calorimetric response, baseline correction, and integration of the calorimetric peaks from the phase transitions of DPPC vesicles, were performed using Origin 8.5 software.

### 2.6. Electron Spin Resonance (ESR)

Continuous wave ESR spectroscopy was carried out at room temperature on a Jeol JES-FA200 spectrometer operating at the X-band (9.2 GHz). The amount of DPPC in protein reconstitution used was 1.36 µmoles, supplemented with 0.5 mol% spin-labelled lipids, 1-palmitoyl-2-stearoyl-(5-doxyl)-sn-glycero-3-phosphocholine (5-PCSL) or 1-palmitoyl-2-stearoyl-(16-doxyl)-sn-glycero-3-phosphocholine (16-PCSL), and protein in a lipid/protein molar ratio of 50. At the final step of the reconstitution protocol, the lipid samples were resuspended in 20 µL of buffer (HEPES 25 mM, NaCl 150 mM, pH 7.4). The proteoliposome solutions containing GRASPs mixed with the spin-labeled DPPC liposomes were drawn into capillary tubes. All ESR spectra were recorded with the following experimental parameters: center field of 3360 G, time constant of 100 ms, field range of 100 G, microwave frequency of 9.2 GHz, modulation frequency of 100 kHz, modulation amplitude of 2.0 G, and microwave power of 10 mW.

ESR spectrum simulations were performed using the Multicomponent program developed by Altenbach and Budil^35^ available at https://www.biochemistry.ucla.edu/Faculty/Hubbell/software.html, which is based directly on the code described by Schneider and Freed^36^. NLSL simulations with MultiComponent allowed us to obtain parameters related to the dynamics, including the rotational diffusion tensor R_⊥_ and the lipid ordering S_0_. The parameters used for the magnetic tensors **g** and **A** (hyperfine interaction) in the fitting of the 5-PCSL labelled DPPC vesicles were: g_xx_ = 2.0089; g_yy_ = 2.0058; g_zz_ = 2.0020; A_xx_ = 4.90 G; A_yy_ = 4.90 G; A_zz_ = 34.70 G^37^, while in the case of 16-PCSL: g_xx_ = 2.0089; g_yy_ = 2.0058; g_zz_ = 2.0021; A_xx_ = 4.90 G; A_yy_ = 4.90 G; A_zz_ = 33.00 G^38^. Information on the spin probe dynamics in the two regions of the lipid chain (5-PCSL and 16-PCSL) was obtained using the parameters R_⊥_, c20, and c22, and, specifically for 16-PCSL, the W_2_ component of the Lorentzian linewidth broadening tensor W. It is important to note that the g and A tensors were kept fixed, whereas R⊥, c20, c22, and W_2_ were varied.

### 2.7. Large Unilamellar Vesicles (LUVs) preparation

The proteoliposomes prepared by the reconstitution protocol were used to prepare unilamellar vesicles for fluorescence experiments. An appropriate volume of DPPC, solubilized in chloroform, was supplemented with 0.5 mol% of either 1-palmitoyl-2-{12-[(7-nitro-2-1,3-benzoxadiazol-4-yl)amino]dodecanoyl}-sn-glycero-3-phosphocholine (hereafter referred to as NBD-C12-PC) or 1,2-dipalmitoyl-sn-glycero-3-phosphoethanolamine-N-(7-nitro-2-1,3-benzoxadiazol-4-yl) (ammonium salt) (hereafter referred to as NBD-PE). The solvent was evaporated, and the resulting lipid film was reconstituted according to the protocol. After lipid resuspension with a detergent-free buffer, Large Unilamellar Vesicles (LUVs) were prepared by extruding the proteoliposome solution with a 100 nm pore polycarbonate membrane from Whatman (Schleicher & Schuel).

### 2.8. Fluorescence Spectroscopy

Fluorescence measurements were performed using a Hitachi F-7000 spectrofluorimeter (Hitachi High-Tech Corporation, Tokyo, Japan) equipped with a 150 W xenon arc lamp. The excitation wavelength was set to 380 nm, with a scan speed of 1200 nm/min, 5-nm excitation and emission slits, a response time of 0.5 s, and a PTM voltage of 700 V. The measurements were performed at 25 °C, and temperature control was achieved using a F25 ultra-thermostatic water bath (JULABO USA, Inc., Pennsylvania, USA), with a sample equilibration time of 3 min.

### 2.9. Giant Unilamellar Vesicles (GUVs) preparation

Giant unilamellar vesicles (GUVs) were produced via electroformation^39^. DPPC, POPC, and Biotinyl-PE stocks were prepared in a 9:1 (vol/vol) CHCl_3_:MeOH mixture. LAURDAN (6-Dodecanoyl-2-Dimethylaminonaphthalene) stocks were prepared in DMSO (5 mM). GUVs used for hyperspectral imaging were made of DPPC containing 0.8 mol% LAURDAN and 0.1 mmol% Biotinyl-PE. For the 3D reconstructions, 0.25 mol% of L-α-Phosphatidylethanolamine-N-(lissamine rhodamine B sulfonyl) was used instead of LAURDAN. A 10 µL volume of a chloroform solution containing 300 µM of lipid was deposited on the surfaces of two conductive glasses (coated with fluorine tin oxide), which were placed with their conductive sides facing each other and separated by a 2-mm thick Teflon frame. The osmolality of the glucose and sucrose solutions used for GUV preparation was carefully adjusted using a cryoscopic osmometer (Osmomat 030, Gonotec, Berlin, Germany). This chamber was filled with a 200 mOsm/kg sucrose solution and connected to a 2 V AC generator with a sinusoidal wave frequency of 10 Hz for 2 hours, followed by a step using 1 Hz for 15 minutes. The vesicle solution was diluted 7 times in a 200 mOsm/kg glucose solution. The glass coating for GUV imaging was prepared according to the protocol described by Moens et al.^40^. Imaging chambers were filled with a 1 mg/mL biotin-labelled BSA solution (Sigma-Aldrich, product number A8549) for 5 min, followed by a 10 min incubation with 5 µg of albumin (Sigma-Aldrich, product number A9275). Pretreatment with Biotin-BSA and avidin immobilizes the Biotinyl-PE-labelled GUVs, forming a sandwich-like configuration with a Biotin-BSA coating on the bottom of the glass slides that interacts with avidin to immobilize the GUVs. After each step of imaging chamber pretreatment, excess protein was washed with a 200 mOsm/kg glucose solution by pipetting. GUVs were added to the coated imaging chambers, and images were taken 15 minutes after protein addition.

### 2.10. Hyperspectral Imaging

Hyperspectral images were acquired using a confocal Zeiss LSM 780 microscope equipped with a 40×, 1.3 NA oil-immersion objective (Carl Zeiss, Germany). A laser wavelength of 405 nm was used for LAURDAN excitation. Image acquisition was performed with a 512 × 512-pixel frame and a 138-nm pixel size. For hyperspectral imaging, the lambda mode (xyλ) of the Zeiss LSM 780 was used, sequentially measuring 32 channels with a bandwidth of 8.94 nm from 410 to 696 nm. The images were processed by the SimFCS software developed at the Laboratory of Fluorescence Dynamics, which is available on the webpage (https://www.lfd.uci.edu/globals/). Because we used a fluorescently labelled protein in the LAURDAN spectral change experiments, we employed a 3-component analysis to separate the protein and lipid signals that are mixed in a single pixel. Spectral phasor analysis used a Fourier transform of the fluorescence emission spectra, from which we calculated two parameters, G and S (as shown in the equations below), representing the real and imaginary components of the transform.

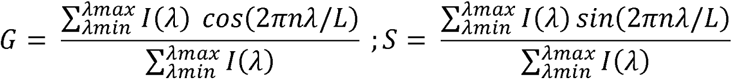

In the equations above, I(λ) is the intensity in each step of spectral measurement, L is the spectrum range (λ_max_ – λ_min_), and n is the harmonic number (=1 in this case). The spectral phasor plot comprises the two parameters G and S plotted in a four-quadrant polar plot. In this case, the wavelength range spans 32 channels from 410 nm to 696 nm, and the spectrum is recorded at each pixel of the hyperspectral image using the Zeiss LSM 780 detector.

### 2.11. Fluorescence imaging of GUVs in contact with myr-GRASPs

Myr-GRASPs were fluorescently labeled with Alexa Fluor 546 NHS Ester (Thermo Fisher, Waltham, MA, United States). The dye, solubilized in ultrapure DMSO (10 mM), was mixed with the purified protein at a dye:protein molar ratio of 1:1, followed by constant agitation at room temperature for 2 hours. The reaction was performed in the dark, followed by separation of the unbound fluorophore from the labeled protein by size-exclusion chromatography using a Hiprep Desalting 26/10 column (Cytiva Life Sciences). The final protein concentration used for all fluorescence confocal microscopy imaging was 0.2 µM, prepared from a 20 µM stock solution. The same volume of buffer without protein was added to control GUVs.

## 3. Results and discussion

### 3.1. Myristoylation of the human GRASPs and its effects on protein secondary structure

Myristoylation is a lipid modification catalyzed by N-myristoyltransferases^41^ (NMTs) in the presence of myristic acid and coenzyme A (CoA). Because this post-translational modification does not naturally occur in prokaryotic organisms, such as *E. coli*, which were employed in this study for heterologous protein expression, we recapitulated the modification in a non-eukaryotic system by co-expressing NMT and Golgi reassembly and stacking proteins (GRASPs) in the presence of myristic acid^30^ (**Figure 1**A). GRASPs were first recognized as central components of the Golgi apparatus. In humans, the two paralogs, GRASP65 and GRASP55, localize at the Golgi by anchoring to cisternal membranes via an N-terminal myristoylation (**Figure 1**B). The myristoylation protocol was first established for the GRASP domain of GRASP55 (DGRASP55),^30,31^ and has now been extended to the other human GRASPs: GRASP55, GRASP65, and the GRASP65 GRASP domain (DGRASP65). In our previous report^31^, we demonstrated the dimerization of the myristoylated GRASP domain of GRASP55 (myr-DGRASP55) in the presence of detergent, highlighting the requirement for this dimerization to facilitate membrane interaction of the lipidated protein. Here, we aimed to evaluate the effects of myristoylation on protein structure and protein-membrane interactions across all human GRASPs, with a particular focus on the possible role of the SPR domain.

**Figure 1.**
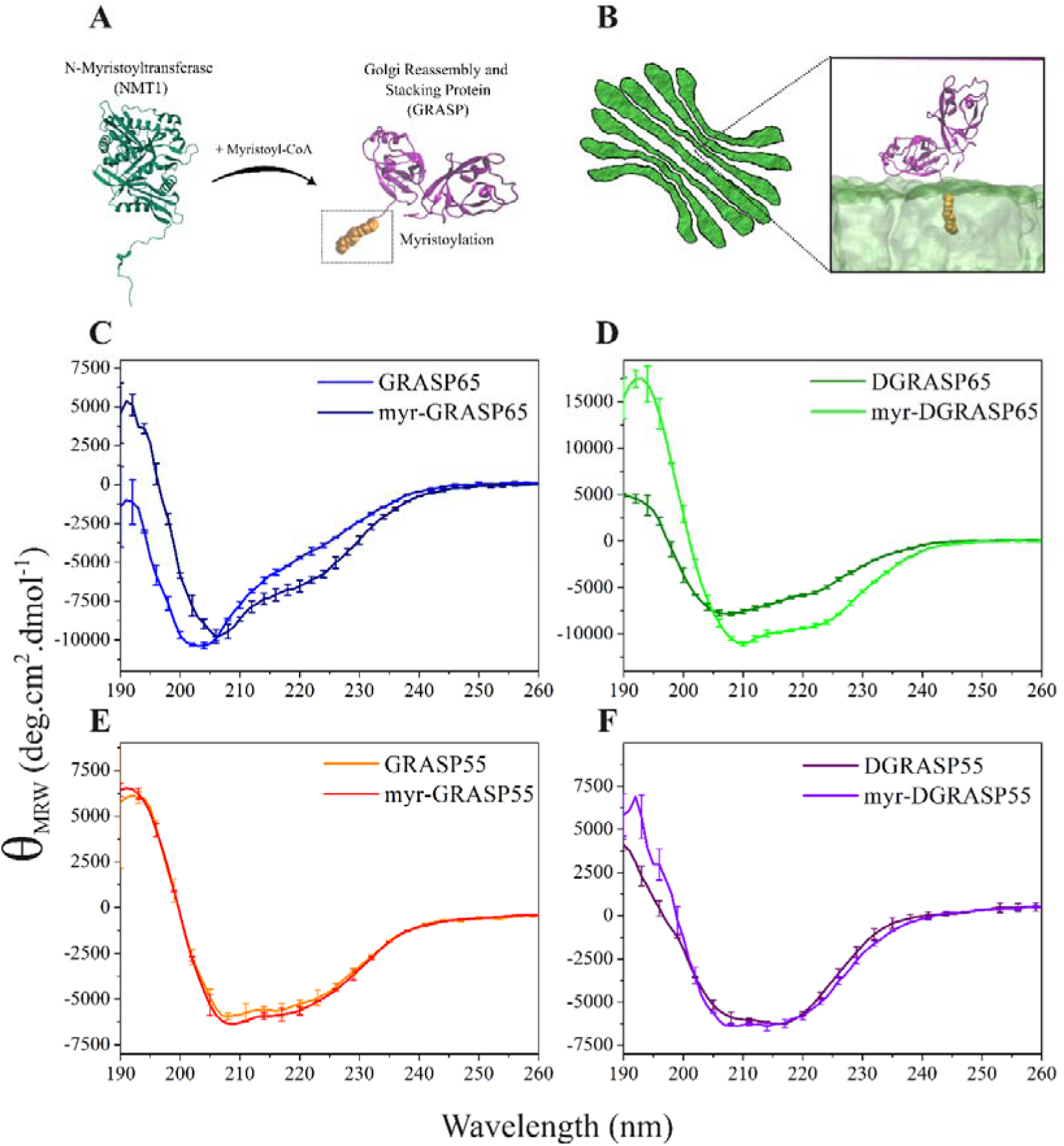
– (A) Myristoylation mechanism catalyzed by NMT1 in the presence of GRASPs and myristoyl-CoA. (B) Human GRASPs are anchored to the Golgi apparatus via N-terminal myristoylation. The figure shows myristoyl-anchored DGRASP55 in lipid membranes, previously obtained from all-atom molecular dynamics simulations^30^. Figure not drawn to scale. The Far-UV CD spectra of human GRASPs in their myristoylated and non-myristoylated forms are shown in (C) GRASP65; (D) DGRASP65; (E) GRASP55, and (F) DGRASP55.

The successfully myristoylated, expressed, and purified human GRASPs (Figures S1 and S2) showed differences in their solubility in the presence and absence of detergent in the purification buffer. The non-myristoylated versions, as expected from previously published data^33,34,42^, were soluble in a free detergent buffer and served as controls to assess structural changes in the myristoylated versions. To check any effect of myristoylation on the overall GRASP structures, we characterized the protein secondary structure by far-UV circular dichroism (CD) spectroscopy (**Figure 1**). The spectra showed no significant changes for myr-GRASP55 and myr-DGRASP55 (**Figure 1**E-F), while myr-GRASP65 (**Figure 1**C) and myr-DGRASP65 (**Figure 1**D) presented pronounced differences caused by myristoylation.

The structural content was estimated by deconvolution of the spectra using the Dichroweb server. The results are shown in Table S1 and the reconstructed spectra in Figure S3. The changes in the secondary structure of GRASP65 caused by myristoylation were particularly pronounced, especially in its GRASP domain. While the helix content of GRASP65 was increased by only 4% by myristoylation, myr-DGRASP65 showed an increase of 17% in its α-helix content and a decrease of 10% in the β-sheet.

### 3.2. HsGRASPs reconstitution into lipid vesicles

We developed a reconstitution protocol for myristoylated GRASPs (myr-GRASPs) and their domains (myr-DGRASPs) to investigate these proteins under conditions mimicking cellular membranes. In summary, we followed the steps widely reported in the literature for the membrane reconstitution of proteins^43^, which involve protein purification in a detergent-containing buffer, followed by detergent removal via hydrophobic adsorption onto polystyrene beads (Bio-Beads) in the presence of liposomes^44^.

The reconstitution assay steps are summarized in **Error! Reference source not found.**A. A molar amount of lipids, 50-fold higher than the protein to be reconstituted, was solubilized in chloroform and dried using N_2_ gas, followed by 2 hours under vacuum to remove any chloroform remnant, forming a lipid film (Step I). After hydrating the lipid film using the purified protein solution in buffer containing DDM (Step II), an isotropic solution of mixed lipids, protein, and detergent micelles was formed by sonicating and vortexing (Step III). The detergent was then slowly removed by adding Bio-Beads under overnight agitation (Step IV), thereby promoting spontaneous association of proteins and phospholipid membranes. Because the adsorbent beads have a small diameter, we found it easier to separate them from the proteoliposome solution using a 200 μL pipette fitted with a 10 μL tip. By positioning the tip close to the bottom of the tube, we prevented beads from being aspirated (Step V). The resulting proteoliposome solution, free of detergent and Bio-Beads, was transferred into centrifuge microtubes (Step VI). Non-anchored proteins were then removed by centrifugation, leaving the reconstituted protein in the pellet fraction (Step VII). Proteoliposome pellets were resuspended in buffer for use in subsequent experiments (Step VIII). Non-reconstituted proteins in the supernatant were quantified by absorbance measurements, enabling calculation of the amount of protein associated with the liposomes (Error! Reference source not found.B). The results in **Figure 2**B show our reconstitution protocol successfully produced proteoliposomes for subsequent investigation.

**Figure 2.**
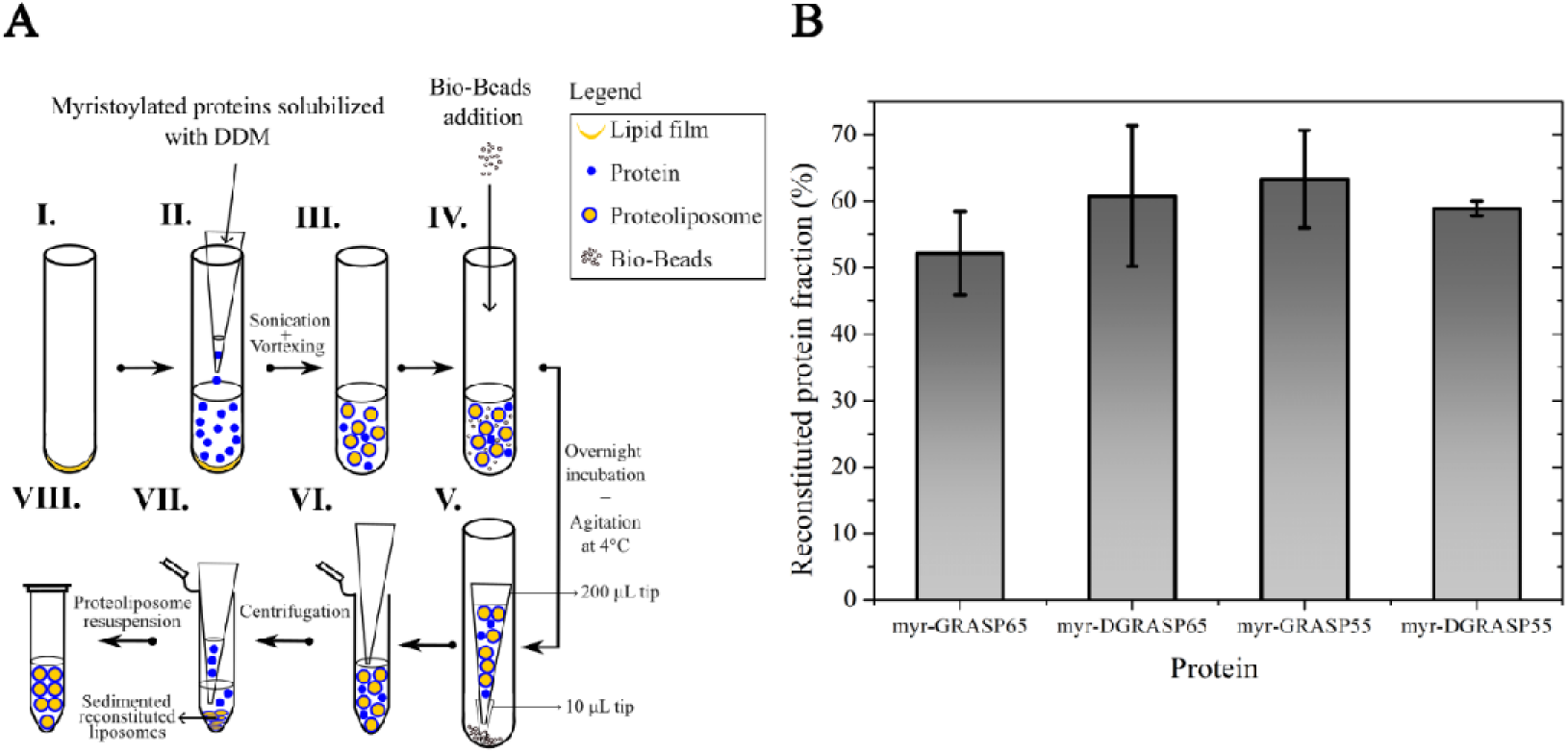
– (A) Reconstitution protocol for myristoylated GRASPs: I) Lipid film formation after chloroform evaporation; II) Lipid film hydration with purified protein solution in DDM buffer; III) Lipid resuspension by ultrasonic bath and vortexing; IV) Bio-Beads addition followed by overnight incubation; V) Proteoliposome solution removal and VI) separated from the beads; VII) Proteoliposome sedimentation and separation of non-reconstituted protein fraction after proteoliposome centrifugation; VIII) Resuspension of the proteoliposome solution in buffer for subsequent experimental use. (B) Protein reconstituted fraction calculated by measuring supernatant concentration after step VII.

### 3.3. Differential Scanning Calorimetry (DSC)

We also investigated the interaction between myristoylated GRASPs and zwitterionic lipid bilayers by examining lipid phase transitions. **Figure 3** shows the thermograms of pure DPPC multilamellar vesicles in the presence and absence of the anchored proteins. Upon heating, pure DPPC exhibited two calorimetric peaks: the pretransition, characterized by a broad and less energetic transition from the tilted gel phase (L_β’_) to the rippled gel phase (P_β’_), and the main phase transition, representing the conversion to the liquid-crystalline phase (L_β_)^45^. In our experiments, the pre– and main transitions were centered at T_P_ = 35.4 °C and T_M_ = 41.2 °C, which agree with the expected values^46^.

**Figure 3.**
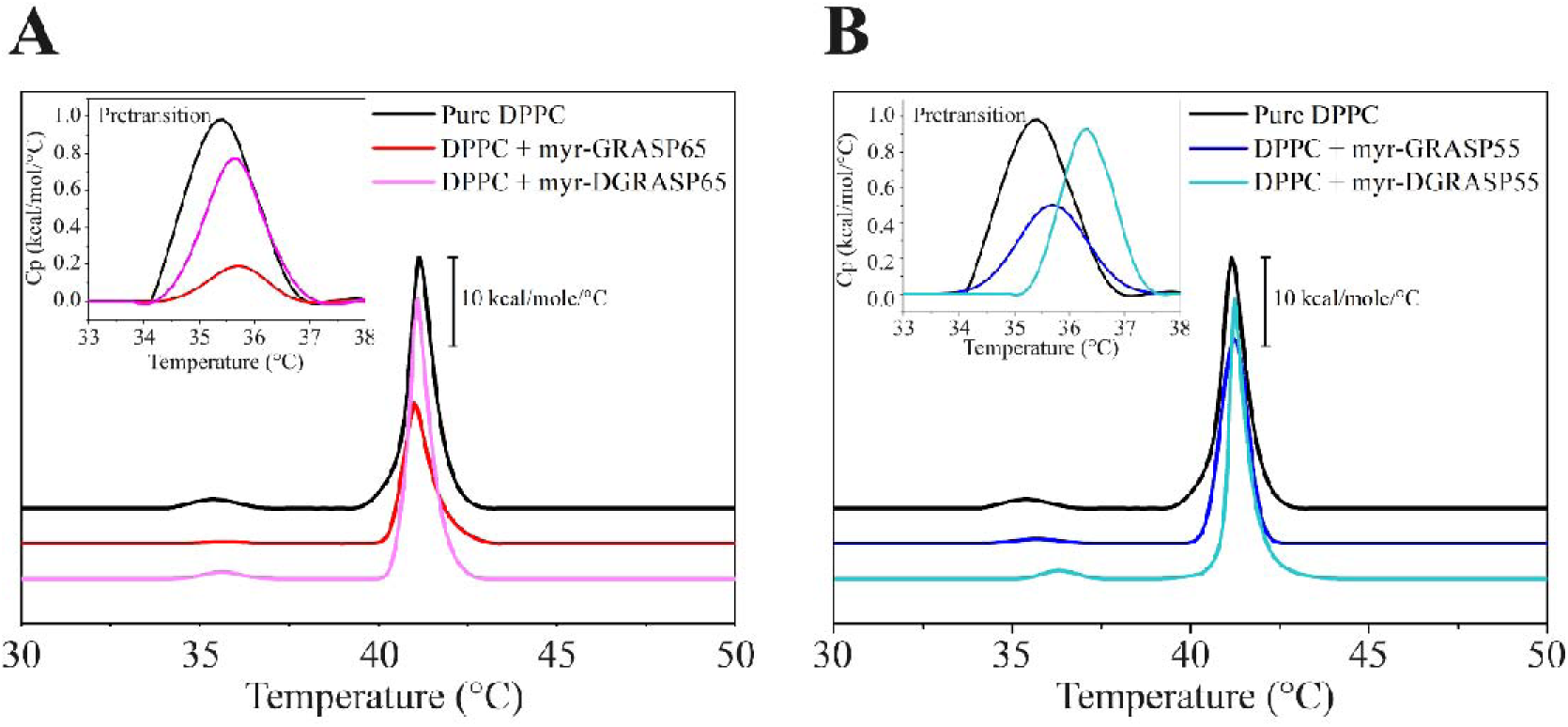
– Heat capacity profiles of DPPC multilamellar vesicles in the presence of reconstituted (A) myr-GRASP65 and myr-DGRASP65 and (B) myr-GRASP55 and myr-DGRASP55. The detailed effects of reconstituted GRASPs on the lipid pretransition are shown in the insets.

The reconstituted GRASPs decreased the enthalpy variation of the main and pretransition compared to the control thermograms (Table S3), with myr-GRASP65 (Figure 3A) having the most significant effect, followed by myr-GRASP55 (Figure 3B), suggesting that the disordered SPR domain makes a substantial contribution to this decrease. The GRASP domains also decreased the enthalpy variations. The slight increase in the pretransition temperature observed in all cases suggests a modest stabilization of the tilted gel (L_β′_) phase relative to the rippled gel (P_β′_) phase. The pretransition in DPPC is highly sensitive to interfacial packing and headgroup orientation, and its upward shift indicates a local increase in headgroup order or a reduction in hydration at the membrane surface. Akin to the pretransition temperatures, the main transition temperatures were not significantly affected by the reconstituted proteins.

The DSC analyses revealed that the myristoylated GRASPs modulate the thermotropic behavior of DPPC bilayers by decreasing the enthalpic variations and cooperativities of both the pre– and main phase transitions, while leaving their transition temperatures unchanged. These findings indicate that GRASP–membrane interactions perturb the lipid packing locally rather than globally. The insertion of the myristoyl moiety into the bilayer likely disrupts interactions between adjacent acyl chains, whereas the disordered SPR domain may interact dynamically with the lipid headgroups, increasing surface heterogeneity and reducing the collective nature of the transition. The preservation of transition temperatures suggests that these perturbations do not significantly alter the average bilayer order or thickness but instead reduce the number of lipids undergoing cooperative melting.

### 3.4. ESR and NLSL simulations

GRASPs are peripheral membrane proteins that interact with lipid membranes through lipidations^30,31^, or in some cases, mediated by an amphipathic helix and a N-terminal acetylation^47^. We have previously described a myristoylation protocol for investigating GRASP/lipid interactions^30^ and demonstrated that myristoylated GRASP55 can dimerize when anchored to model membranes^31^. However, aspects regarding the protein’s influence on lipid organization remained unclear. To characterize the lipid membranes with attached GRASPs, we measured ESR spectra using spin-labeled phospholipids. The ESR spectrum is highly sensitive to the spin label ordering and dynamics^48–50^. Additionally, we performed NLSL simulations, which enabled us to properly quantify the lipid membrane dynamics. We used two spin probes: one with the nitroxide radical at the acyl chain carbon 5 (5-PCSL, **Figure 4**A), and the other in a position closer to the end of the lipid chain at carbon 16 (16-PCSL, **Figure 4**B). The environment experienced by 5-PCSL is closer to the membrane surface, while 16-PCSL probes the middle of the lipid bilayer. This strategy enables us to assess the extent to which myristoyl-anchored GRASPs can impact the lipid microenvironment. The experimental and calculated spectra obtained from the simulations using the MultiComponent program are shown in **Figure 4**C.

**Figure 4.**
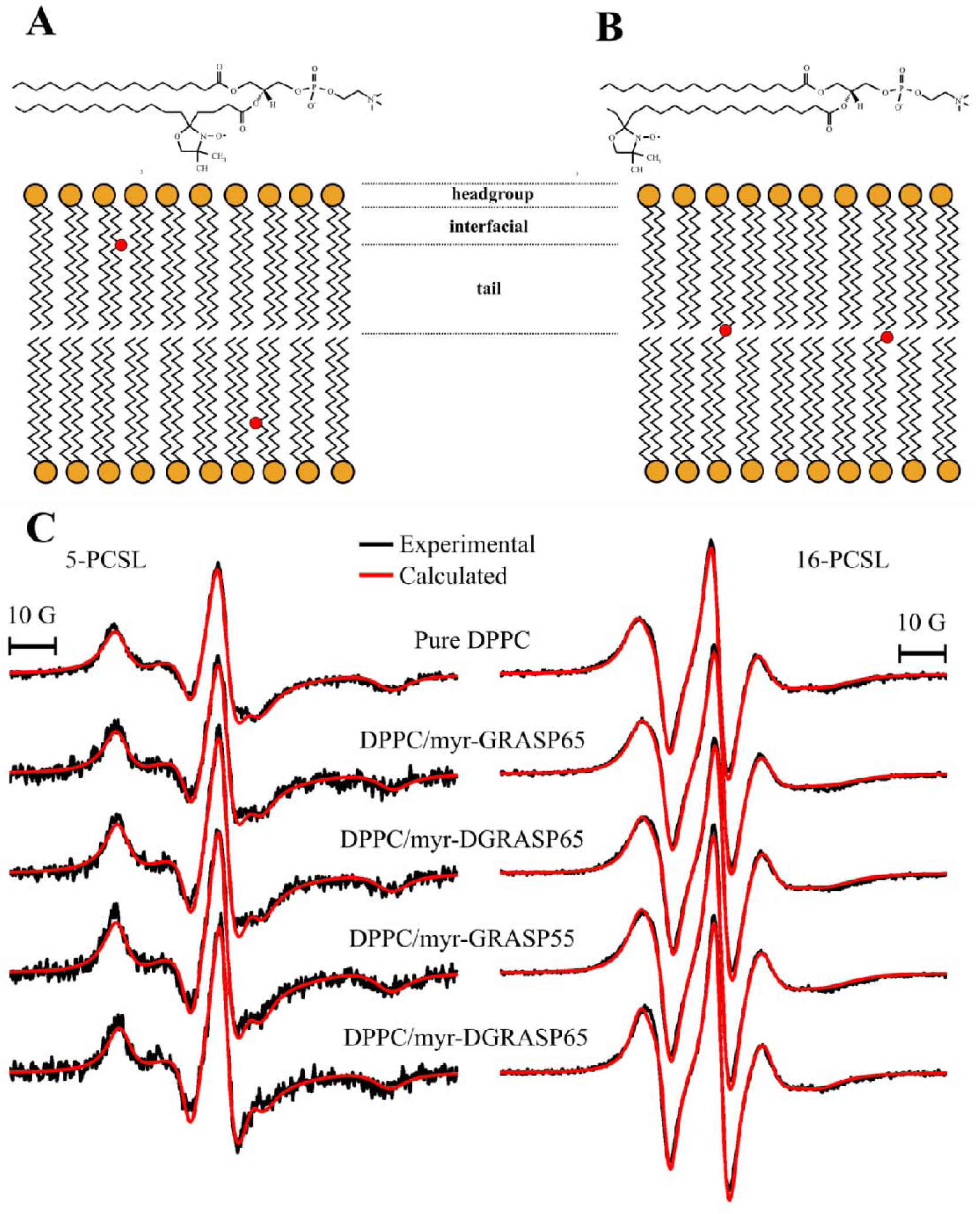
– Structures of (A) 1-palmitoyl-2-stearoyl-(5-doxyl)-sn-glycero-3-phosphocholine (5-PCSL) and (B) 1-palmitoyl-2-stearoyl-(16-doxyl)-sn-glycero-3-phosphocholine (16-PCSL). The membrane bilayer schematic shows the location of the spin probes (red dot) in a DPPC lipid environment. (C) 5-PCSL and 16-PCSL ESR spectra at room temperature of DPPC vesicles in the absence and the presence of myristoylated GRASPs in their full-length form and only the GRASP domain. The red lines represent the best theoretical fits obtained from NLSL simulations. Total width: 100 G.

The overall lineshapes of the 5– and 16-PCSL in pure DPPC bilayers are markedly different^49^, reflecting the different microenvironments they experience near the membrane surface (5-PCSL) versus in the middle of the bilayer (16-PCSL). The broader 5-PCSL spectrum reflects the restricted motion of a highly packed region of the membrane bilayer^51^. For the control DPPC vesicles containing 5-PCSL, the spectral simulation yielded an R_1_ value of 0.049 × 10^9^ s^-1^, corresponding to a correlation time τ_c_ of 3.40 ns, and an order parameter (S_0_) of 0.28. The reconstituted 5-PCSL-DPPC vesicles containing reconstituted proteins yielded a bilayer environment in which the spin probe exhibited faster motion in the presence of the proteins. We observed this slightly enhanced membrane fluidity for the vesicles containing the anchored myr-GRASP65, where the rotational diffusion rate R_1_ calculated from the simulated spectrum was 0.055 × 10^9^ s^-1^ (τ_c_ = 3.03 ns) and S_0_ of 0.33. When considering only the GRASP domain of the membrane-attached GRASP65, myr-DGRASP65 yielded an R_1_ of 0.069 × 10^9^ s^-1^ (τ_c_ = 2.40 ns) and an S_0_ of 0.40. This effect was more pronounced for myr-DGRASP55, where the simulations yielded an R_1_ of 0.089 × 10^9^ s^-1^ (τ_c_ of 1.87 ns) and an S_0_ of 0.48. For the full-length GRASP55 instead, the R_1_ of 0.050 × 10^9^ s^-1^ (τ_c_ of 3.33 ns) and an S_0_ of 0.31 presented a weaker effect in increasing the lipid bilayer dynamics when compared to myr-DGRASP55. Overall, the membrane becomes slightly more fluid in the presence of myristoylated proteins, with the greatest effect observed with myr-DGRASP55. Furthermore, our results suggest that the SPR domain imposes a slight decrease in the spin probe dynamics at the 5^th^ position of the acyl chain.

We also assessed the nitroxide microenvironment further along the lipid acyl chain using the spin probe 16-PCSL, which is located closer to the bilayer center, where chain mobility is naturally higher. In this case, the NLSL simulations of the control DPPC resulted in a R_1_ 0.138 × 10^9^ s^-1^, τ_c_ 1.20 ns, and S_0_ 0.13. Those parameters did not show significant changes in the presence of the lipidated GRASPs (Table S2).

### 3.5. Fluorescence spectroscopy

Fluorescence spectroscopy has been a significant tool for studying lipid membrane structures by using extrinsic membrane probes that can be covalently attached to specific regions of the lipid molecule, which can then render information about structural and dynamical properties of the surroundings where the fluorescent probe is linked to, similarly to what was done in the previous section using electron spin resonance (ESR). A popular family of fluorophores used to study membrane dynamics is the 7-nitrobenz-2-oxa-1,3-diazol-4-yl (NBD), which can be linked to various lipids and helps study different processes, including membrane phase separation using resonance energy transfer and similar methodologies^52^, lateral membrane mobility using fluorescence recovery after photobleaching (FRAP)^53^, and also the spatial organization of proteins or other molecules in contact with lipid membranes^54^. In our case, we explored the dynamics of NBD in two situations involving GRASP-reconstituted lipid vesicles: the fluorophore attached to the lipid headgroup (NBD-PE, **Figure 5**A) and the lipid tail, specifically at the 12^th^ carbon position (NBD-C12-PC, **Figure 5**B). In the DPPC lipid bilayer, NBD-PE positions the fluorophore near the membrane surface at the polar headgroup region, whereas NBD-C12-PC localizes it deeper within the hydrophobic core, around the bilayer midplane (**Figure 5**C).

**Figure 5.**
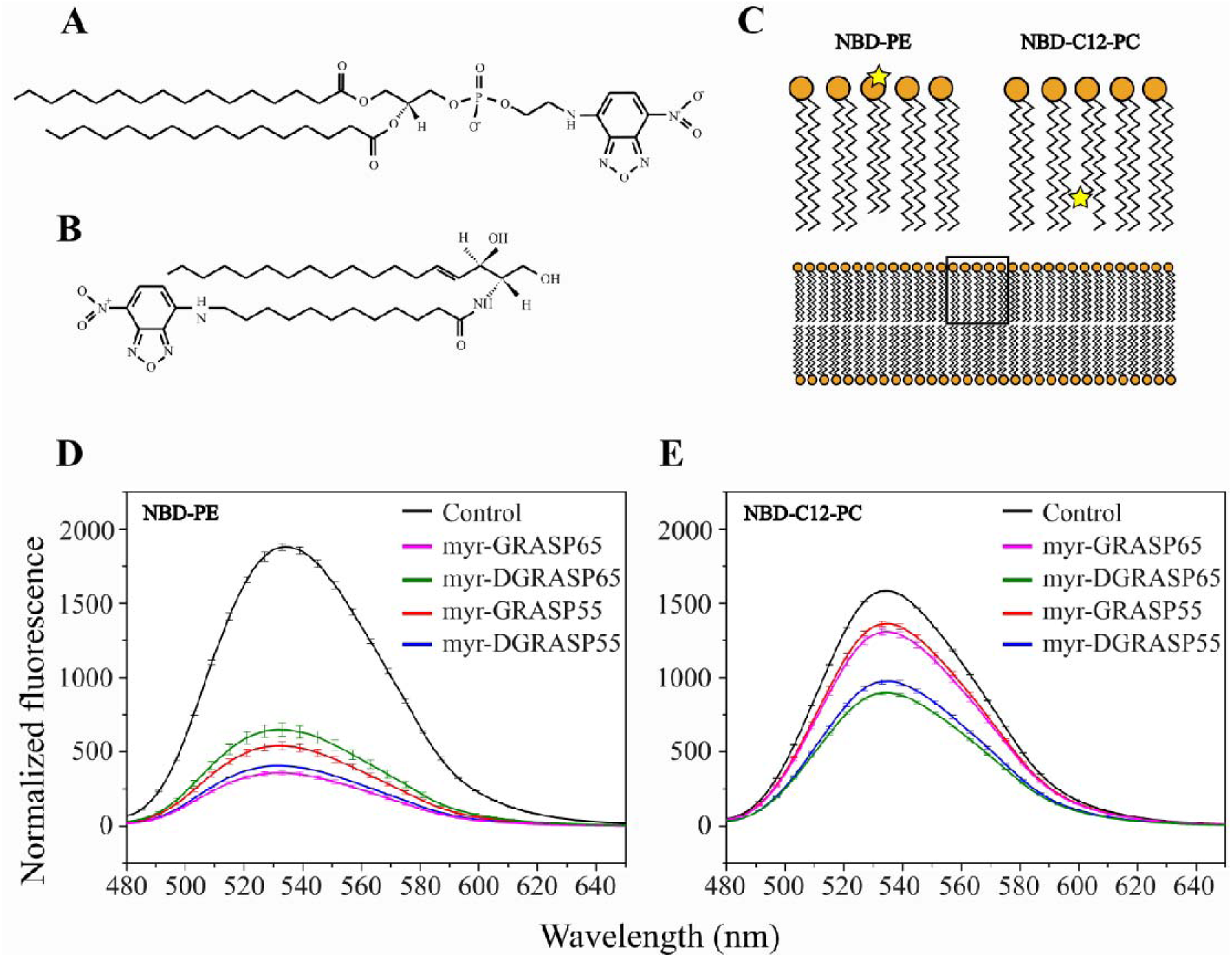
– Molecular structures of (A) NBD-PE and (B) NBD-C12-PC. (C) Schematic diagram of a lipid bilayer, where the yellow star shows the region where the fluorophore is located, in the lipid headgroup in the case of NBD-PE, and at position 12 of one of the lipid carbon chains, in the case of NBD-C12-PC. Fluorescence spectra of 7-nitrobenz-2-oxa-1,3-diazol-4-yl (NBD) positioned at (D) the lipid headgroup (NBD-PE) and (E) the 12^th^ carbon at the acyl chain (NBD-C12-PC), in the presence of the myristoyl-anchored GRASPs.

We measured the fluorescence spectra of NBD in DPPC liposomes, both in the absence and in the presence of reconstituted GRASPs. A first notable characteristic is that the spectral intensity is reduced in the presence of the anchored proteins, even after absorbance normalization. For NBD-PE, a slight change in the emission maximum was also observed in the presence of GRASPs, a ∼ 3 nm blue shift that might indicate changes in the microenvironment polarity at the lipid headgroup region when the protein was membrane-attached (**Figure 5**D). This change, however, was not observed for NBD-C12-PC (**Figure 5**E), which appears to be less affected by the myristoyl-anchored GRASPs, consistent with the 16-PCSL results from ESR experiments. The lower NBD fluorescence intensity observed in both cases likely reflects a decrease in NBD quantum yield associated with changes in the local membrane environment upon protein binding. In particular, protein-induced perturbation of DPPC packing may increase water penetration into the bilayer, thereby enhancing quenching of NBD fluorescence. This effect is expected to be more pronounced for the headgroup-located NBD probe, which is more exposed to the aqueous interface, than for NBD-C12-PC residing deeper in the membrane.

### 3.6. Hyperspectral imaging (HSI)

After characterizing the potential influence of the SPR domain on protein-membrane interactions using various spectroscopic methods and calorimetry, we then investigated these interactions using fluorescence microscopy. To do so, we used giant unilamellar vesicles (GUVs) as membrane models, whose dimensions (between 1-100 µm) were compatible with the imaging method, allowing us to investigate changes promoted by the presence of the proteins.

We performed hyperspectral imaging experiments using GUVs containing LAURDAN as a fluorescent probe. LAURDAN is a fluorophore from the DAN family, first synthesized by Gregorio Weber and collaborators^55^ for monitoring spatial and temporal polarity and water dipolar relaxation at the probe nano-environment^56^. Due to its higher hydrophobicity when compared to other DAN derivatives, LAURDAN has been used to explore the structure of biological and biomimetic membranes using fluorescence microscopy^57,58^. We investigated the effects of proteins on membrane fluidity using spectral phasor analysis of LAURDAN fluorescence spectrum^59^. By applying the phasor approach, one can detect spectral shifts in LAURDAN-labelled GUVs induced by changes in the membrane microenvironment, even in the presence of other fluorescent species. The phasor is computed over a specific wavelength range, and the phasor coordinates will shift depending on the fluidity of GUVs, as demonstrated in the phasor plot calculated from the emission spectra of LAURDAN in the gel (L_β_) and liquid disordered (Ld) phases of the membrane (Figure 6A)^60,61^. Since the protein was also labelled with a fluorescent dye (Alexa546), we performed a 3-component analysis to separate lipid and protein signals, represented by the cyan triangle in Figure 6A. In GUVs containing LAURDAN, the presence of a third fluorescent component at a given pixel, indicated by the blue cursor in Figure 6A, will cause the phasor at that pixel to lie along the line connecting the LAURDAN in the gel phase and the fluorescent protein. We can separate complex spectral mixtures using linear combination rules, thereby characterizing membrane fluidity. **Figure 6**A shows the extent of LAURDAN fluorescence in POPC (green), representing the Ld phase, and DPPC (red), representing the gel phase, combined with Alexa546-labeled protein, forming a theoretical triangle encompassing all possible linear combinations within the cyan triangle. The blue cursor in the phasor plot indicates the fluorescent GRASP signal.

**Figure 6.**
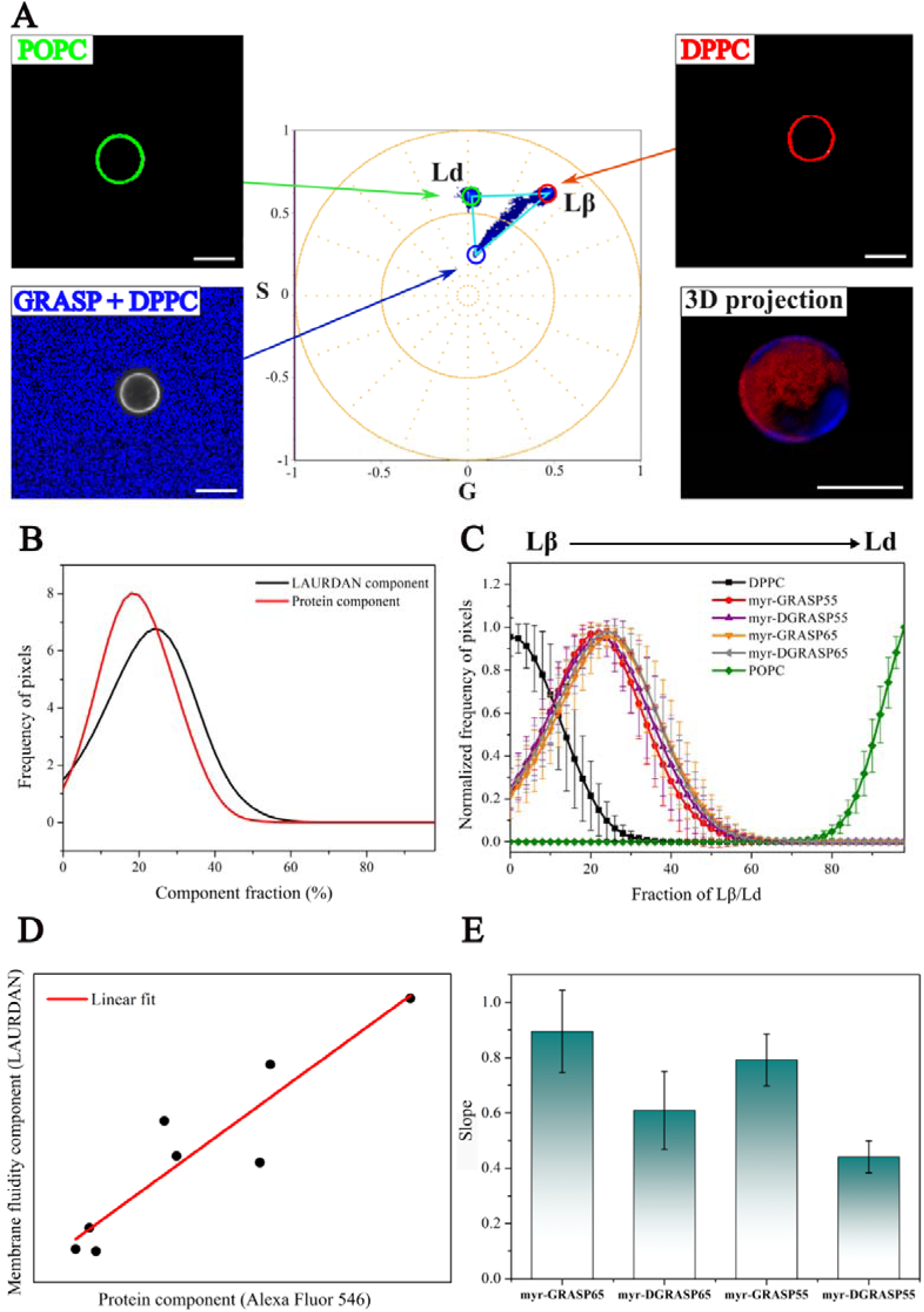
– Three-component analysis describing LAURDAN fluorescence emission in the gel (L_β_) phase in DPPC GUVs, liquid disordered phase (Ld) in POPC vesicles, and the effects of myristoylated GRASPs on DPPC GUVs fluidity. (A) Three-component analysis configuration by defining the emission of LAURDAN in the Lβ (DPPC, red GUV) or Ld (POPC, green GUV) membranes, and the fluorescent signal of GRASPs (colored by the dark blue cursor). (B) Example of the component histograms that are generated for each image. (C) LAURDAN component calculated for the different samples, showing an increase in the average membrane fluidity upon incubation with protein. (D) Representative example of the linear fit analysis performed by integrating each lipid and protein component histogram, assuming a linear dependence between the amount of protein and GUV fluidity. (E) The slope of the fitted lines indicates a stronger influence on the lipid membrane microenvironment caused by the full-length myr-GRASPs. Scale bars = 10 µm.

In images of GUVs incubated with protein, several pixels showed mixed signals from the protein-labeling fluorophore, Alexa 546, and LAURDAN. The 3-component analysis in SimFCS4 enabled separation of the fluorescence emission of two fluorophores in the phasor distribution, yielding a histogram containing two distributions for each phasor component (protein and lipid) in each image (**Figure 6**B). **Figure 6**C shows the LAURDAN component distribution for each sample. The LAURDAN histograms of the protein-incubated GUVs show increased overall water dipolar relaxation and membrane polarity, indicating enhanced membrane fluidity caused by myristoylated GRASPs. To quantify how the protein component affected the displacement of the LAURDAN component in the phasor plots, we integrated the distribution histograms of each component. We compared them with the histograms of control GUVs using the same component analysis. As shown in Figure 6C, the average distribution of the LAURDAN and Alexa546 components in the control sample (black line) shows a limitation of up to 30% in the histogram (Figure S4A); therefore, we used this point as a cutoff to start integrating the histograms of both components for the protein-incubated samples. The integrated areas are shown in Figure S4B. We plotted the integrated areas of the protein and lipid components for each image from each sample to identify a relationship between membrane fluidity and the amount of protein measured in the GUV images. To do so, the protein versus lipid component plots were assumed to behave as the simplest model, a linear behavior between protein amount and membrane fluidity (**Figure 6**D). The linear fits showed that the full-length myr-GRASPs had more influence in increasing the water dipolar relaxation and polarity experienced by the LAURDAN-labelled GUVs (**Figure 6**E). This result is noteworthy, indicating that the SPR domain influences membrane dynamics.

## 4. Discussion

GRASPs are present in several branches of the eukaryotic tree of life^62^. The interaction involving lipid membranes and the representatives of this family has been shown to occur in different ways. In some earlier eukaryotes, as in the case of the GRASP65 homologue 1 (Grh1) in *Saccharomyces cerevisiae*, this affinity with membranes is mediated by the presence of an amphipathic helix and an acetylation at the N-terminus^47^. *Plasmodium falciparum* possesses two GRASP isoforms that interact with membranes differently: one exemplar with a structure similar to fungal GRASPs, and another with a myristoylated N-terminus, comparable to higher eukaryotes^63^. In Metazoans, the evolutionary process led to the emergence of two GRASP paralogues, gorasp1 and gorasp2, which encode the myristoylated proteins GRASP65 and GRASP55, respectively^4,30,62^

The primary goal of this study was to establish an efficient protein-membrane reconstitution protocol for human GRASPs and evaluate how these proteins can affect lipid membranes. We used the previously described DGRASP55 myristoylation protocol^30^ to myristoylate full-length GRASP55 and GRASP65, as well as the GRASP domain of GRASP65 (DGRASP65). Since we reported that myr-DGRASP55 was soluble in the absence of detergent, we purified all myristoylated GRASPs in free-detergent solution, yielding highly soluble myr-GRASP65 and myr-GRASP55 and poorly soluble myr-DGRASP65. This difference between the highly soluble myr-GRASP65 and the lower solubility observed for myr-DGRASP65 suggests that myr-GRASP65 may also possess a myristoyl-switch mechanism, similar to what we observed for myr-DGRASP55^31^, regulated by its SPR domain. Our circular dichroism spectra support this hypothesis, suggesting that the overall secondary structures were affected by myristoylation (**Figure 1**), with myr-DGRASP65 and myr-GRASP65 being more affected.

Our DSC data showed that the pretransition of DPPC liposomes was affected by the presence of reconstituted GRASPs. The transition temperatures were less affected by the presence of the proteins (Table S3). On the other hand, we observed a substantial reduction in the enthalpy variation of the pre– and main transitions for the full-length GRASPs proteoliposomes (**Figure 3**A). The cooperativity of the lipid transition can be directly determined from the sharpness of the transition peak, as reflected in the temperature width at half-height (ΔT_1/2_). Here, we observed a more pronounced effect of reduction in DPPC gel-to-liquid crystalline transition cooperativity caused by myr-DGRASPs (**Figure 3**B and Table S3). The result was not observed in our thermograms for the myr-GRASPs. The main transition in lipid bilayers is associated with abrupt conformational changes occurring in the lipid acyl chains^64^. In this sense, the effect of a reduced enthalpy of the main transition induced by the presence of the anchored myr-DGRASPs may indicate a more pronounced influence on the organization of the DPPC hydrocarbon chains.

In the context of the study of myristoyl-anchored GRASPs reported here, ESR and fluorescence spectroscopy provided important insights into the effects of GRASPs on lipid organization in a membrane-mimetic system. The ESR spectra of the two probes yielded different effects, with 5-PCSL revealing decreased membrane order and increased membrane dynamics (**Figure 4**C). Moreover, variations in these properties were more pronounced for the GRASP domains, suggesting that the presence of the SPR domain somehow affects the membrane-binding properties of these proteins. The spectra obtained using the probe 16-PCSL (**Figure 4**D) did not reveal significant changes upon the anchoring of GRASPs, indicating that the changes occur mostly at carbons closer to the headgroup of the lipid chain.

Complementary to ESR, we also investigated the effects of reconstituted proteins on lipid organization using NBD-labeled lipids in different regions of the lipid acyl chain and explored the fluorescence properties of these probes in various environments (**Figure 5**). The probes NBD-PE and NBD-C12-PC showed decreased fluorescence intensity in the presence of anchored GRASPs. The reduction in fluorescence intensity is likely associated with changes in the local membrane environment induced by protein anchoring, such as perturbation of lipid packing and increased water penetration, which are known to decrease the fluorescence quantum yield of NBD (**Figure 5**). The magnitude of NBD-PE fluorescence quenching did not differ significantly among the different constructs, suggesting that this probe at the lipid headgroup is not sufficiently sensitive to discriminate between their effects. In contrast, for NBD-C12-PC, the extent of quenching clearly distinguishes between proteins, with myr-DGRASPs exhibiting a noticeably higher ability to cause NBD fluorescence quenching compared to the other constructs.

Overall, the DSC, ESR, and fluorescence data reveal a consistent, depth-dependent effect of myristoylated GRASPs on membrane organization. The reduction in enthalpy and cooperativity of the DPPC transitions observed by DSC, which was most pronounced for myr-DGRASPs, indicates a global loosening of lipid packing. ESR measurements at the bilayer interface (5-PCSL) show this effect, resulting in decreased order and increased dynamics, whereas the deeper 16-PCSL probe reports minimal changes. Likewise, NBD-C12-PC fluorescence measurements revealed that myr-DGRASPs exert the strongest local perturbations, suggesting that the GRASP domains can penetrate deeper into the lipid bilayer.

The LAURDAN hyperspectral imaging experiments provide direct evidence that myristoylated GRASPs enhanced membrane fluidity in zwitterionic lipid systems (**Figure 6**). The spectral phasor analysis reveals a clear shift of the LAURDAN emission toward higher dipolar relaxation values in GUVs incubated with the full-length myr-GRASPs, indicating an increase in local water penetration and lipid dynamics at the bilayer interface (**Figure 6**C). Together, these findings support a model in which membrane-anchored GRASPs perturb bilayer organization through the combined action of their myristoyl anchor and disordered SPR domain (**Figure 6**E). The myristoyl group likely inserts into the hydrophobic core of the bilayer, introducing packing defects and local curvature stress, while the flexible SPR domain may interact transiently with the membrane surface, enhancing headgroup hydration and lateral mobility. The observed correlation between protein abundance and increased LAURDAN relaxation further suggests a dose-dependent effect, in which greater protein coverage amplifies membrane fluidization. Overall, the fluorescence microscopy data complement the calorimetric and spectroscopic results, demonstrating that GRASPs modulate membrane physical properties by reducing lipid packing order and increasing interfacial hydration, effects that may underlie their role in shaping and organizing Golgi membranes in vivo.

## 5. Concluding remarks

In this study, we report the membrane interactions of human GRASPs modified at their N-termini with a myristoyl group. When reconstituted into liposomes, GRASPs can affect lipid bilayer structure and dynamics, with clear differences observed between the presence and absence of the SPR domain, indicating a role for this domain in regulating the membrane interactions of GRASP65 and GRASP55. Moreover, our findings regarding the effect of GRASPs on membrane fluidity designate an interesting effect of this protein that can be related to the participation in Golgi stacking and vesicular unconventional processes observed *in cell*. To the best of our knowledge, this is the first detailed report on the in vitro effects of myristoylated GRASPs on the dynamic structure of phospholipid model membranes at the molecular and microscopic levels.

## 6. Acknowledgments

The authors gratefully acknowledge the financial support of the National Institute of Science and Technology in Innovative Research in Health Sciences – from Nanotechnology to Artificial Intelligence (INCT PICS) sponsored by Brazil’s National Council for Scientific and Technological Development (CNPq), grant no. 408417/2024-2, Coordination of Superior Level Staff Improvement (CAPES), grant no. 88887.197686/2025-00, and São Paulo Research Foundation (FAPESP), grant no. 2025/26818-7. AJCF and RI thank FAPESP for the partial support via grants 2024/17967-6, 2024/17969-9, 22023/04532-9, and 2021/10795-7, and CNPq via grants 309476/2025-9 and 311831/2021-4. LM and MD were supported by the grants 2020-225439, 2021-240122, and 2022-252604 of the Chan Zuckerberg Initiative DAF, an advised fund of the Silicon Valley Community Foundation, and by FOCEM – Fondo para la Convergencia Estructural del Mercosur (COF 03/11).

