## Supplementary data for "Exploring the effects of Golgi Reassembly and Stacking Proteins in lipid membranes"

^1^Departamento de Física, Faculdade de Filosofia, Ciências e Letras de Ribeirão Preto, Universidade de São Paulo, Avenida Bandeirantes, 3900, 14040-901 Ribeirão Preto, Brazil

^2^Advanced Bioimaging Unit, Institute Pasteur de Montevideo - Universidad de la República, Montevideo, Uruguay

^3^Departamento de Fisiopatología, Hospital de Clínicas, Facultad de Medicina, Universidad de la República, Montevideo, Uruguay

^4^Instituto de Física, Universidade de São Paulo, Rua do Matão, 1371, 05508-090, São Paulo, Brazil

^5^National Institute of Science and Technology in Innovative Research in Health Sciences: from Nanotechnology to Artificial Intelligence, Ribeirão Preto, SP, Brazil

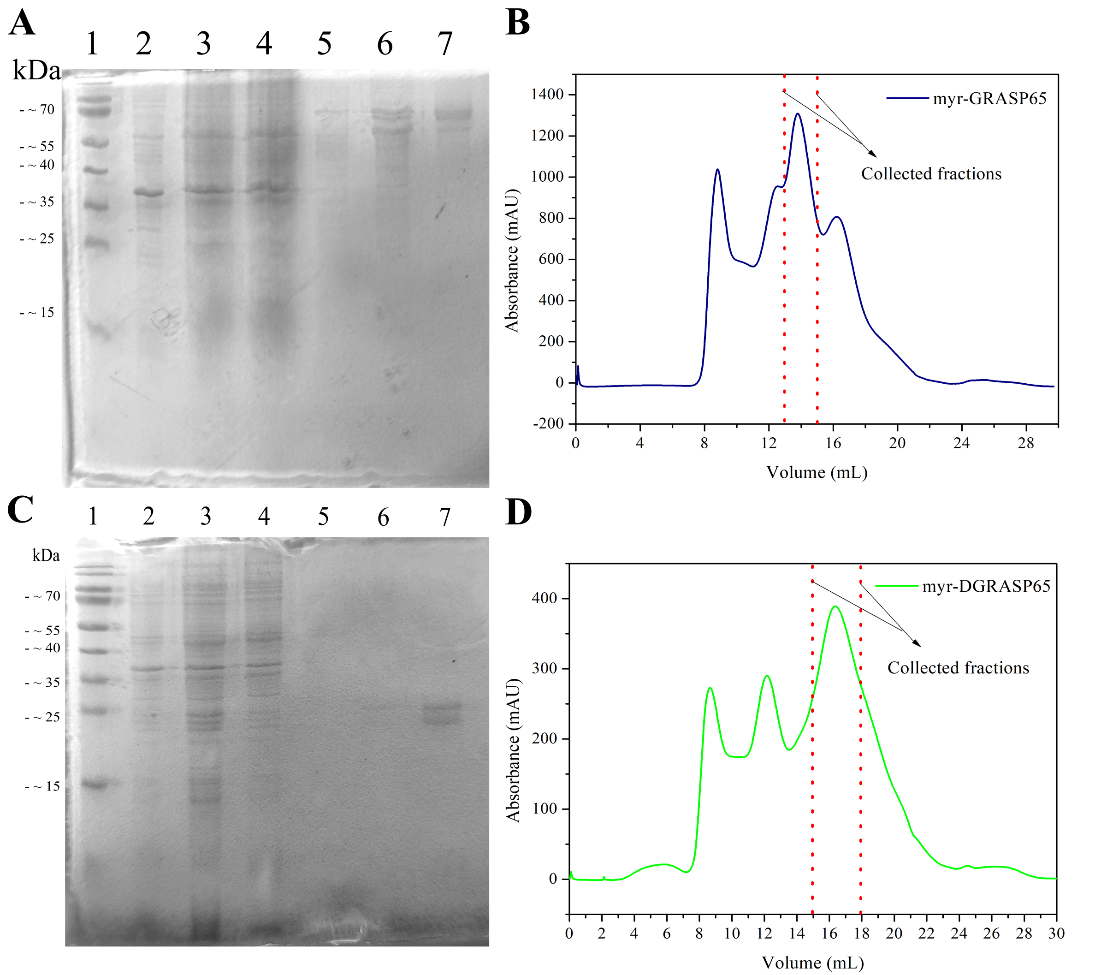

Figure S1 – Purification of myr-GRASP65 and myr-DGRASP65. SDS-PAGE results for (A) myr-GRASP65 and (C) myr-DGRASP65 purification steps: 1 – Protein standard molecular weight ladder; 2 - Cell pellet after lysis and centrifugation; 3 – Supernatant after lysis and centrifugation; 4 – Fraction unbound to the column; 5 – 20 mM imidazole washing; 6 – Protein elution using 300 mM imidazole and 7 – Purified protein fraction collected from the size exclusion chromatography column. Gel filtration chromatograms of (B) myr-GRASP65 and (D) myr-DGRASP65, where the SEC collected fractions are indicated by the red dotted lines.

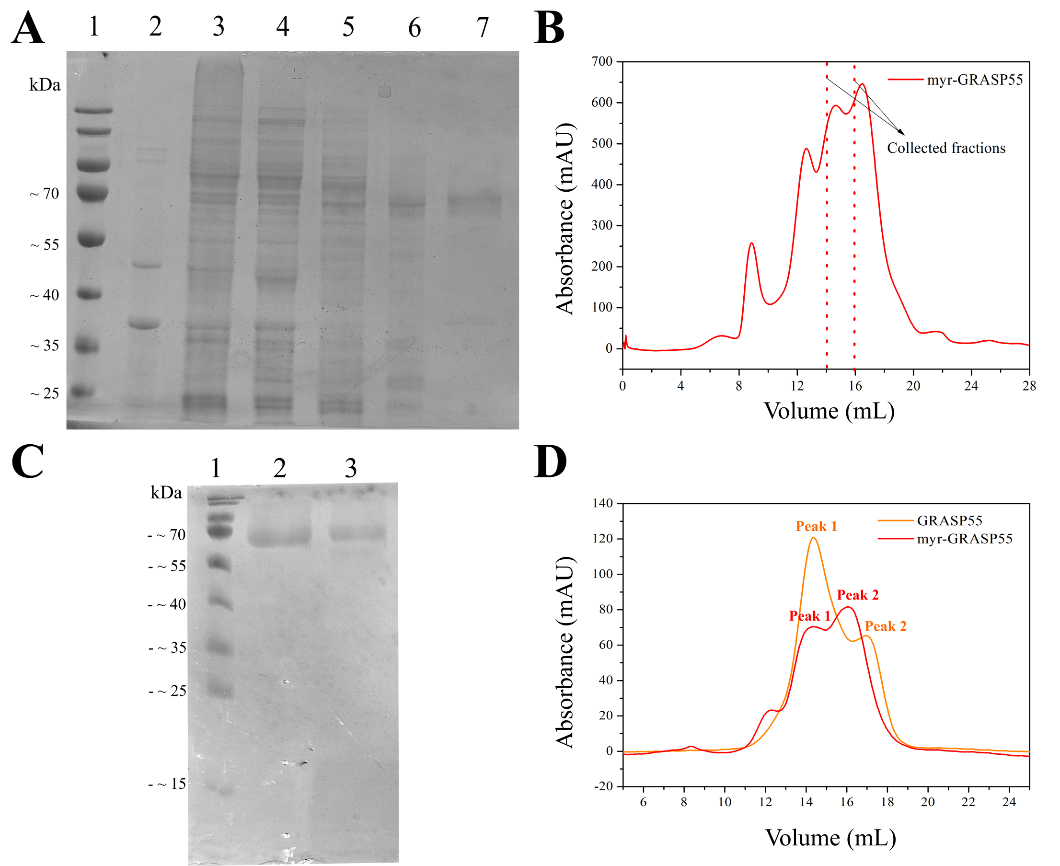

Figure S2 - Purification of myr-GRASP55. SDS-PAGE results for (A) myr-GRASP55 and purification steps: 1 – Protein standard molecular weight ladder; 2 - Cell pellet after lysis and centrifugation; 3 – Supernatant after lysis and centrifugation; 4 – Fraction unbound to the column; 5 – 20 mM imidazole washing; 6 – Protein elution using 300 mM imidazole and 7 – Purified protein fraction collected from the size exclusion chromatography column. The last step of purification chromatogram using gel filtration is presented in (B), where the SEC collected fractions are indicated by the red dotted lines. (C) Comparison between the collected fractions of GRASP55 and myr-GRASP55 depicted in the chromatogram shown in (D). In lane 1 – Protein standard molecular weight ladder; 2 – GRASP55 SEC eluted samples (mixing fractions peak 1 and peak 2) and 3 – myr-GRASP55 eluted samples (mixing fractions peak 1 and peak 2). The graph in D is the reinjection of the ‘peak 1’ previously separated from the SEC column for GRASP55 and myr-GRASP55 results in splitting again in two different peaks.

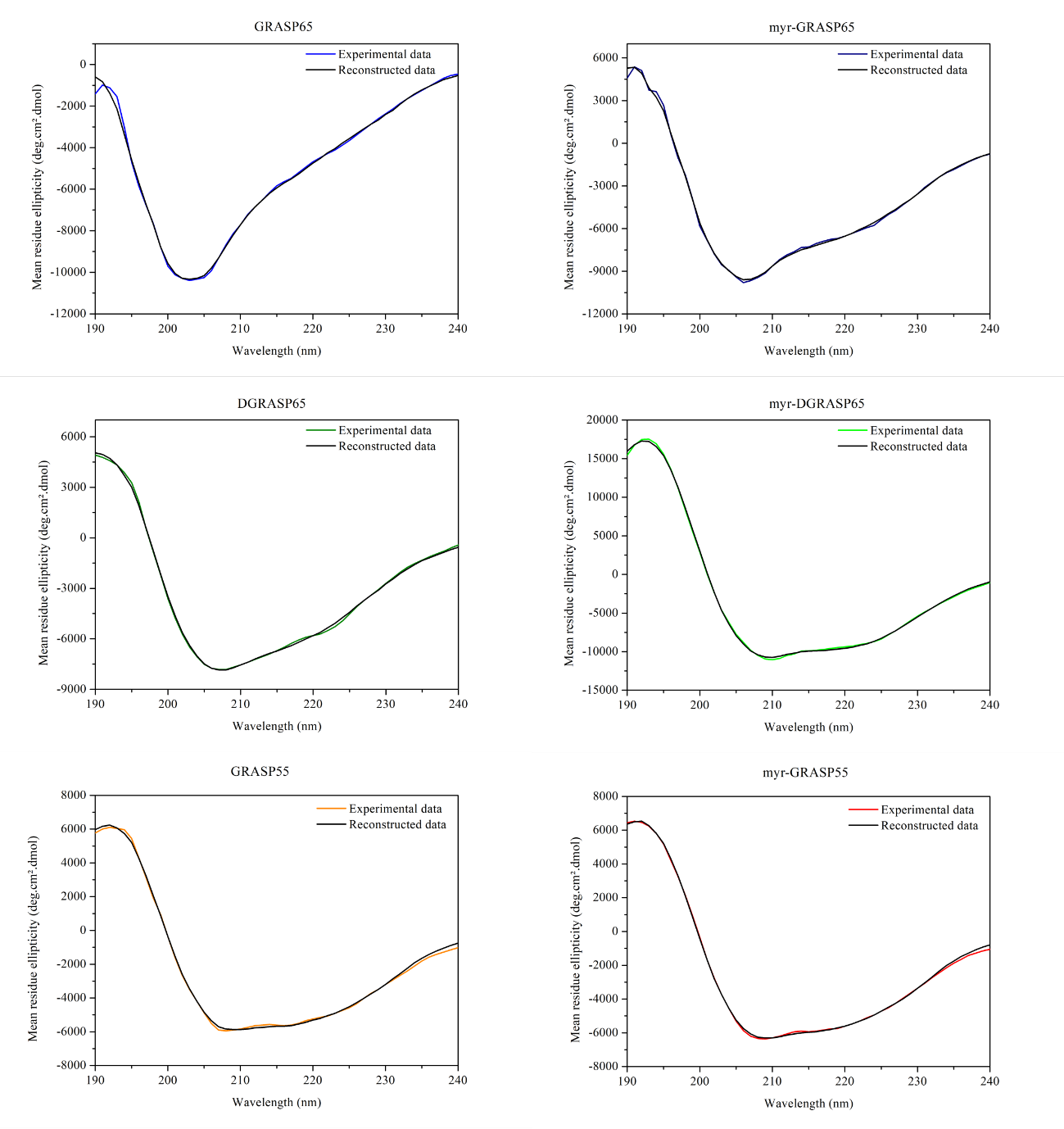
Figure S3 – Deconvolution of the far-UV CD spectra of myristoylated GRASPs using the CDDSSTR method. Theoretical (reconstructed) spectra are shown as black lines.

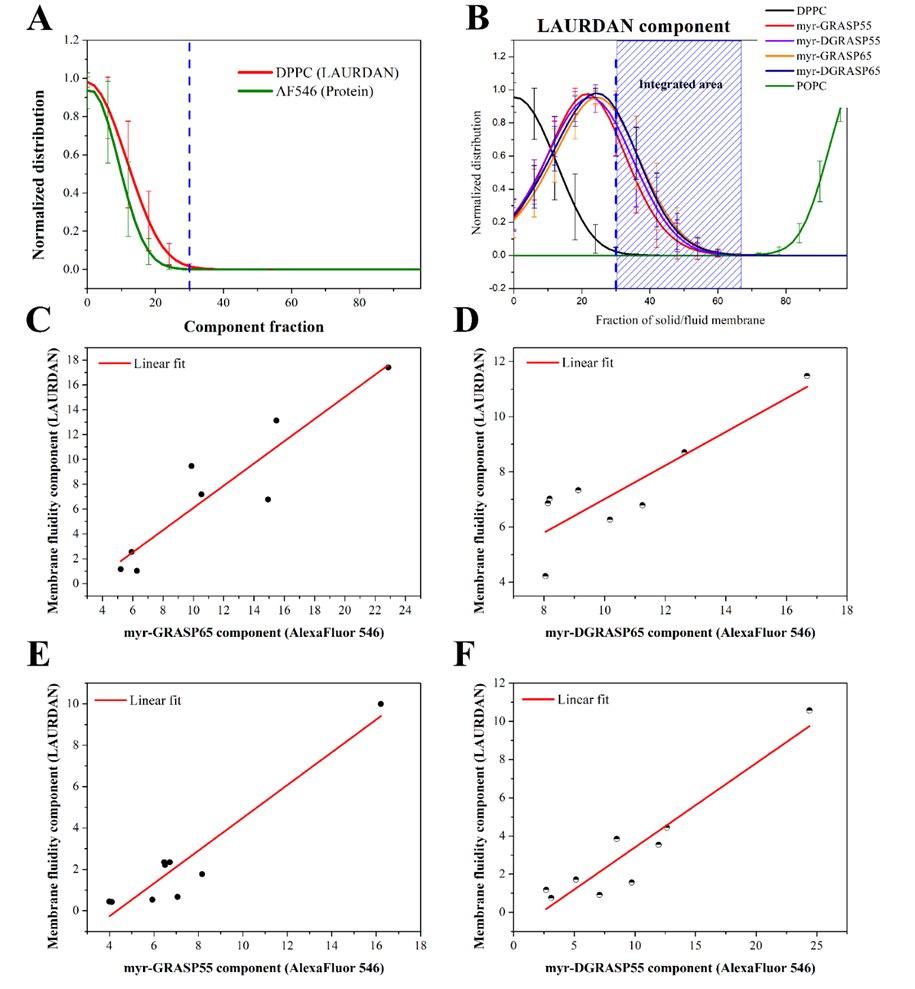
Figure S4 – Three component analysis of lipid and protein components. A) Histograms generated from the 3-component analysis for the control DPPC GUVs, where the blue dashed line indicates the starting point for the integration of the histograms used for the protein-incubated vesicles. B) Resulting distributions of the LAURDAN component obtained for each image for the 4 different versions of myristoylated GRASPs calculated, where the blue dashed area corresponds to the integrated area used in the plotting of protein versus lipid component. Each image analyzed using the three-component analysis generated two histograms that were integrated according to the delimitations shown in A) and B). The linear fits of the integrated histograms are shown for C) myr-GRASP65; D) myr-DGRASP65; E) myr-GRASP55 and F) myr-DGRASP55.

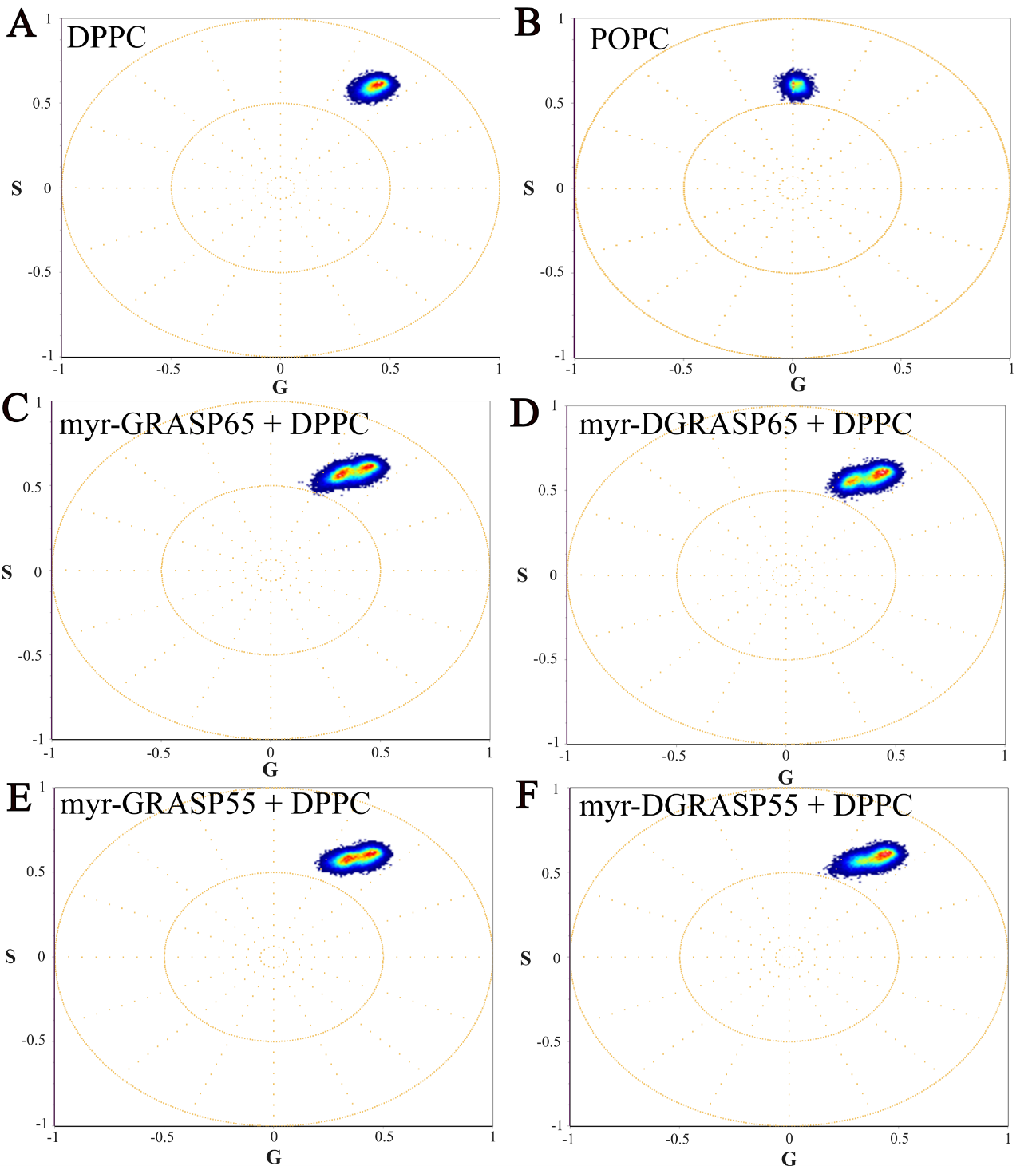

Figure S5 – Phasor plots obtained for each sample considering all spectral images acquired. A) DPPC; B) POPC; The following phasor plots consist on DPPC incubated with C) myr-GRASP65; D) myr-DGRASP65; E) myr-GRASP55 and F) myr-DGRASP55.

Table S1 – Secondary structure content calculated from the deconvolution of the CD spectra using the CDSSTR method in the Dichroweb webserver. The deconvolution data of DGRASP55 and myr-DGRASP55 was obtained from our previous report^1^.

| **Protein** | **Helix (%)** | **Strands (%)** | **Others (%)** |
| --- | --- | --- | --- |
| GRASP65 | 12 | 29 | 59 |
| Myr-GRASP65 | 16 | 28 | 56 |
| DGRASP65 | 14 | 32 | 54 |
| Myr-DGRASP65 | 31 | 22 | 47 |
| GRASP55 | 13 | 32 | 55 |
| Myr-GRASP55 | 14 | 31 | 55 |
| DGRASP55 | 9 | 34 | 57 |
| Myr-DGRASP55 | 14 | 26 | 60 |

Table S2 – Rotational diffusional rate (R_1_), Correlation time (τ_c_) and order parameter (S_0_) values obtained from the simulations of spin-labeled DPPC liposomes in the presence of anchored myr-GRASPs. The lipid/protein molar ratio was estimated by calculating the protein concentration using absorbance (280 nm) in the supernatant fraction after protein reconstitution into liposomes.

| Sample | 5-PCSL | | | 16-PCSL | | |  |
| --- | --- | --- | --- | --- | --- | --- | --- |
|  | R_1_ (×10^9^ s^-1^) | τ_c_ (ns) | S_0_ | R_1_ (×10^9^ s^-1^) | τ_c_ (ns) | S_0_ | Lipid/protein molar ratio |
| Control | 0.058 | 2.83 | 0.269 | 0.170 | 0.99 | 0.119 | - |
| Myr-GRASP65 | 0.061 | 2.72 | 0.300 | 0.151 | 1.10 | 0.107 | 95.8 |
| Myr-DGRASP65 | 0.070 | 2.37 | 0.329 | 0.178 | 0.94 | 0.124 | 82.3 |
| Myr-GRASP55 | 0.061 | 2.75 | 0.310 | 0.166 | 1.01 | 0.104 | 79.1 |
| Myr-DGRASP55 | 0.078 | 2.12 | 0.375 | 0.209 | 0.99 | 0.119 | 84.9 |

Table S3 - Thermodynamic parameters of the pre- and main transition of DPPC, obtained from the analysis of the thermograms of the reconstituted myristoylated GRASPs.

| Pretransition | | | Main transition | | | |
| --- | --- | --- | --- | --- | --- | --- |
| Sample | T_p_ (°C) | ∆H_p_ (kcal/mol) | T_M_ (°C) | ∆H_M_ (kcal/mol) | ∆T_1/2_ (°C) | ∆S (cal/[mol.K]) |
| Pure DPPC | 35.4 | 1.45 | 41.24 | 31.9 | 0.85 | 0.77 |
| DPPC + myr-GRASP65 | 35.7 | 0.23 | 41.00 | 15.8 | 0.84 | 0.38 |
| DPPC + myr-DGRASP65 | 35.6 | 1.01 | 41.10 | 25.0 | 0.66 | 0.61 |
| DPPC + myr-GRASP55 | 35.7 | 0.77 | 41.19 | 21.7 | 0.89 | 0.53 |
| DPPC + myr-DGRASP55 | 36.3 | 1.06 | 41.24 | 22.2 | 0.52 | 0.54 |

Table S4 – Slope values calculated from the linear fits obtained for each protein sample incubated with DPPC GUVs shown in Figure S4.

| **Protein** | **Slope** | **Standard Error** |
| --- | --- | --- |
| myr-GRASP65 | 0.8944 | 0.1496 |
| myr-DGRASP65 | 0.6091 | 0.1496 |
| myr-GRASP55 | 0.7918 | 0.0942 |
| myr-DGRASP55 | 0.4408 | 0.0578 |

1.	Kava, E., Garbelotti, C.V., Lopes, J.L.S. & Costa-Filho, A.J. Myristoylated GRASP55 dimerizes in the presence of model membranes - PubMed. *Journal of Biomolecular Structure & Dynamics* **43**, 6530-6451 (2024).
